# Evaluation of the Peacock Eye Disease Management (PedMan) System in Highly Susceptible Olive Cultivar and Using the System for Addressing Key Biological Questions

**DOI:** 10.64898/2026.06.02.729514

**Authors:** Yotam Gilat, David Ygzao, Dani Shtienberg, David Ezra

**Affiliations:** Department of Plant Pathology and Weed Research, Institute of Plant Protection, Agricultural Research Organization - Volcani Institute, Rishon LeZion 7528809, Israel; The Robert H. Smith Faculty of Agriculture, Food and Environment, The Hebrew University of Jerusalem, Rehovot 7610001, Israel

**Author notes:** Corresponding author: DE. Authors with equal contribution (Yotam Gilat, David Ygzao).

**Keywords:** control, decision support system, epidemiology, *Venturia oleaginea*

## Abstract

Peacock eye disease, caused by *Venturia oleaginea*, is a major foliar disease of olive (*Olea europaea*) in Mediterranean regions, resulting in defoliation, reduced tree vigor, and yield losses. Effective disease management relies on precise fungicide timing. In this study, we evaluated a decision support system (DSS) named PedMan (Peacock eye disease Manager), designed to predict infection events and optimize fungicide application timing in highly susceptible olive cultivars based on rainfall and temperature conditions. PedMan recommendations were examined in seven independent grove experiments conducted during the 2023/24 and 2024/25 growing seasons in commercial olive orchards in Israel. In addition, simulation analyses were performed using weather data from 11 meteorological stations representing diverse climatic regions. Field evaluation showed that fungicide applications timed according to PedMan recommendation significantly reduced leaf abscission by approximately 60% compared with untreated controls. Applications made contrary to system recommendations did not improve disease control, confirming the reliability of both positive and negative recommendations. Multi-season analyses indicated cumulative disease suppression, with up to 85% reduction in leaf abscission after three consecutive years of correctly timed applications. The system was implemented by olive growers in 2025/6 with applicable success. Simulation results showed that most infection alerts occurred in autumn and early winter. Across all regions and seasons, PedMan recommended 0- 3 fungicide applications per season, comparable to or fewer than conventional spray programs. These findings demonstrate that PedMan is a robust, field-validated DSS that improves fungicide timing, enhances disease control efficiency, and supports sustainable management of peacock eye disease without increasing spray instances under Mediterranean conditions.

## Introduction

Olive peacock eye disease, also known as olive leaf spot, is one of the most widespread and economically important foliar diseases of olive (*Olea europaea* L.) worldwide (Buonaurio et al. 2023; Graniti 1993; Obanor et al. 2008; Saad and Masri 1978). The disease is caused by the hemibiotrophic fungus *Venturia oleaginea* (syn. *Spilocaea oleaginea*; *Fusicladium oleagineum*) and is characterized by circular, dark green to black lesions surrounded by chlorotic halos, resembling the pattern on a peacock’s tail feathers. Under favorable environmental conditions, severe infections cause premature defoliation, reducing photosynthetic capacity and tree vigor and potentially resulting in substantial yield losses (Graniti 1993). Repeated defoliation may also increase susceptibility to other biotic and abiotic stresses and contribute to long-term declines in productivity (Graniti 1993; Issa et al. 2019; Miller 1949; Obanor et al. 2008, 2011). Disease development is favored by humid, mild conditions, with infection closely associated with rainfall, leaf wetness duration, and moderate temperatures, making Mediterranean regions particularly conducive to disease development (Obanor et al. 2005; Trapero and Blanco 2010; Viruega et al. 2011).

Olive production is an important agricultural sector in Israel, where the cultivar Souri accounts for more than 80% of the cultivated area and is highly susceptible to peacock eye disease (Barazani et al. 2014). Recent studies provided new insights into the epidemiology of *V. oleaginea* in Israel, identifying two distinct periods of infection events during each growing season: autumn-early winter and spring-early summer (Ygzao et al. 2026c). Two corresponding disease-development periods were identified. The first occurs in autumn-early winter and results from infections established during the spring of the previous season, whereas the second occurs in late spring-early summer and results from infections occurring during the autumn-early winter of the current season. These observations are broadly consistent with previous studies conducted in Mediterranean countries (Graniti 1993; Trapero and Blanco 2010; Viruega et al. 2011, 2013). Based on these findings, Ygzao et al. (2026c) concluded that *V. oleaginea* is monocyclic within a growing season, completing only one disease cycle per season. Consequently, effective fungicide applications immediately before or after the autumn-early winter infection period may provide season-long control, with additional applications offering little benefit. Because the disease is polyetic, fungicide applications before or after the spring-early summer infection period may also reduce over summering inoculum and disease severity in the following season (Ygzao et al. 2026c). Field experiments in commercial groves further supported these conclusions, showing that one or a few fungicide applications in the autumn were sufficient to effectively suppress disease development during the same season (Ygzao et al. 2026b), consistent with previous observations by Obanor et al. (2008) and Viruega and Trapero (1999).

As for other plant pathogens, *V. oleaginea* development is strongly influenced by weather conditions (Buonaurio et al. 2023; Issa et al. 2017). Accordingly, fungicide applications should ideally be timed according to environmental conditions rather than fixed calendar schedules. Several empirical and semi-empirical models have been developed to predict disease development from weather variables such as rainfall, temperature, relative humidity, and leaf wetness duration. These forecast or warning systems range from empirical models based on field observations to mechanistic models that simulate pathogen development and infection processes over time (Rhimini et al. 2020; Romero et al. 2018; Roubal et al. 2013; Thomidis et al. 2021; Viruega and Trapero 1999). By analyzing observed or forecast weather data, such models identify periods of high infection risk, commonly referred to as infection alerts. Forecast models provide the basis for decision support systems (DSSs), which integrate environmental and epidemiological information to optimize disease-management decisions (Shtienberg 2013). DSSs may combine forecast-model outputs with real-time or forecast weather data, host phenology, cultivar susceptibility, and historical disease pressure to generate site-specific infection-risk assessments. For example, the Novaterra DSS simulates infection events using hourly temperature and leaf wetness data to generate an infection-risk index for Mediterranean regions (NOVATERRA Project 2024). The main practical advantage of forecast models and DSSs is their ability to optimize fungicide timing by targeting periods immediately before, during, or after infection events rather than relying on predefined spray schedules.

Despite their potential, the practical use of forecast models and DSSs for managing peacock eye disease remains limited (Thomidis et al. 2021), largely because of the lack of independent field examination. To our knowledge, no published studies have implemented grove-scale fungicide programs in which forecast-model or DSS predictions were used to determine spray timing. The only relevant report concerns the Novaterra DSS, for which project results indicated that groves managed according to DSS recommendations had disease incidence similar to or lower than conventionally treated groves despite fewer fungicide applications (NOVATERRA Project 2024).

Based on our previous studies (Ygzao et al. 2026a, b, c), we developed a DSS named PedMan (Peacock eye disease Manager). PedMan integrates environmental information to identify infection events and recommend optimal fungicide application timing. An infection event is defined here as a period with conditions favorable for conidial liberation, dispersal, germination, and penetration. Rainfall is the primary driver of *V. oleaginea* conidial liberation and dispersal, as raindrop impact dislodges conidia from infectious lesions and facilitates their spread to nearby foliage (Buonaurio et al. 2023; Graniti 1993; Trapero and Blanco 2010; Viruega et al. 2013). Accordingly, rainfall is considered a prerequisite for an infection event in PedMan; in its absence, conidia are not effectively dispersed and an infection event is not expected.

Following dispersal, conidia are exposed to solar radiation, desiccation, and fluctuating temperatures. *V. oleaginea* conidia are highly sensitive after detachment from infected leaves and generally lose viability within hours to a few days, although survival may be prolonged under humid and shaded conditions (Agosteo and Schena 2011; Viruega et al. 2013). Once conidia have been dispersed by rainfall, leaf wetness duration and temperature are the main environmental factors governing infection (Buonaurio et al. 2023; Guechi and Girre 1994). Under the Mediterranean conditions prevailing in Israel, leaf wetness results from both rainfall and dew. Rainfall therefore contributes to both conidial dispersal and the wetness required for germination and infection, linking these two processes. Dew is generally deposited during the night and dries in the early morning, under Mediterranean climate; consequently, temperatures during rainfall and during the subsequent nocturnal and early-morning wet periods are critical for infection.

Accurate measurement of leaf wetness duration is challenging because leaf wetness is not a direct atmospheric variable but the result of interactions among relative humidity, air and leaf temperature, solar radiation, wind, and leaf-surface properties. Consequently, different sensors may produce different readings under identical environmental conditions (Sentelhas et al. 2007). Spatial variability within tree canopies further limits the representativeness of single-point measurements, while artificial sensors do not fully reproduce the physical and chemical properties of natural leaf surfaces (Madeira et al. 2002). Differences in sensor calibration and wetness thresholds among studies also limit comparability. These limitations make direct leaf-wetness measurements difficult to standardize and support the use of integrated approaches combining measurements, physical modeling, and expert interpretation (Magarey et al. 2005).

Given these difficulties, and because rainfall can provide the wetness required for conidial germination, PedMan predicts infection events using rainfall and temperature criteria derived from our previous studies (Ygzao et al. 2026a, b, c). For highly susceptible olive cultivars, an infection alert is issued when rainfall reaches at least 15 mm in one day or accumulates to ≥15 mm over several consecutive days, and the average minimum temperature during rainy days is 12–18 °C (Ezra et al. 2026). Infection alerts can be generated from either observed or forecast weather data. PedMan recommends applying a registered fungicide within 14 days before or after the infection alert. The minimum interval between consecutive applications is 14 days. When harvest is imminent, spray timing and fungicide selection must also account for pre-harvest intervals to avoid unacceptable fungicide residues on fruit.

Before commercial implementation, models must be rigorously evaluated to ensure adequate accuracy (Thomidis et al. 2021). Evaluation comprises two complementary stages: verification and validation. Verification assesses whether the model has been correctly implemented and operates as intended, including correct algorithm coding, data processing, and output generation. Validation, in contrast, determines whether the model accurately represents disease dynamics and provides reliable predictions under independent field conditions. After defining the PedMan criteria, weather data from the groves used for model development were analyzed to determine whether the system generated infection alerts at appropriate times. These analyses indicated that PedMan predictions were accurate (Ezra et al. 2026; Gilat et al. 2026), supporting successful verification. The next step was to validate PedMan under commercial-grove conditions that were independent of model development.

Accordingly, the first objective of this study was to evaluate whether fungicide applications timed according to PedMan recommendations effectively suppress peacock eye disease in commercial olive groves. This objective was addressed during the 2023/24 and 2024/25 growing seasons. In the subsequent 2025/26 season, PedMan was implemented by two olive growers in commercial groves. Because the system’s recommendations are based on the prediction of infection events, its performance under commercial field conditions also provided an opportunity to evaluate the validity of the underlying epidemiological assumptions. Therefore, the second objective was to use the validated system to investigate various aspects of peacock eye disease development across the major olive- growing regions of Israel.

## Materials and Methods

### Evaluating the performance of PedMan in grove experiments under natural conditions

#### Experimental sites

PedMan recommendations were implemented in seven experiments carried out during the 2023/4 and 2024/5 growing seasons in four sites. Experiments 1 and 4 were carried out in both seasons in Kfar Ben Nun (31.864651 N; 34.952042 E); Experiments 2 and 5 in Haruvit (31.751613 N; 34.856304 E); Experiments 3 and 6 in Kfar Kish (32.667857 N; 35.465225 E). Experiment 7 was carried out in 2024/5 in another grove in Kfar Ben Nun (31.859262 N; 34.951702 E). Kfar Ben Nun and Haruvit are located in the Inner Plains and Kfar Kish in the Lower Galilee Mountains region. The experiments were carried out in commercial groves of cv. Souri severely infected by peacock eye disease. In all sites, the trees were more than 30 years old, spaced 6 to 7 m apart within rows and 7 m apart between rows. During the experiments, cultural practices including irrigation, fertilization, pruning, insect and weed control, and harvesting were performed by the growers according to the recommendations published by the Israeli extension service.

#### Experimental procedures and treatments

Before the initiation of experiments in Kfar Ben Nun and Haruvit the groves had not been treated with fungicides against *V. oleaginea*. The site of Kfar Kish served us during the previous two growing seasons (2021/2 and 2022/3) to examine aspects related to the timing of fungicide sprays (Ygzao et al., 2026b). Most of the trees included in experiments 3 and 6 of the current report were not sprayed with fungicides against *V. oleaginea* in the previous growing seasons. However, as will be outlined below, there were also few trees that were sprayed in 2022/3. Weather data used in PedMan (rain quantity and minimum temperature) were obtained from the web site of the Israel Meteorological service and the Ministry of Agriculture and Food Security (www.agrometeorology.co.il). Nakhshon, Revadim and Serin Height weather stations are located less than 2 km from the experimental sites in Kfar Ben Nun, Haruvit and Kfar Kish, respectively.

All experiments included two basic treatments: **(i) Control-Control**: trees in that treatment were not sprayed with fungicides against *V. oleaginea* in the previous and in the current growing seasons; **(ii) Control-PedMan**: trees in that treatment were not sprayed with fungicides in the previous growing season but were sprayed according to PedMan recommendations in the current growing season. Experiments 1, 2 and 3 continued in a second growing season (2024/5) and consisted of an additional treatment **- (iii): PedMan-PedMan**: trees in that treatment were sprayed with fungicides according to PedMan recommendations two growing seasons in a row. Experiment 6 consisted of another treatment **- (iv): PedMan-Control**: trees in that treatment were sprayed with fungicides according to PedMan recommendations in the 2023/4 growing season but remained untreated in the 2024/5 growing season. Sprays in experiments 1 and 2 were applied on 21 Nov and 11 Dec 2023 (for the 2023/4 growing season); sprays in experiment 3 were applied on 22 Nov, 12 Dec 2023 and 16 Apr 2024 (for the 2023/4 growing season). In experiment 6 PedMan alerted that infection event had occurred in 18-20 Nov 2024 (for the 2024/5 growing season). Nevertheless, since the grower intended to harvest the crop shortly, spraying was postponed until after harvesting and it was applied 12 days later, on 2 Dec 2024. The season-long rains, average minimum temperature during the rain events and the timing of fungicide applications in the different experiments are presented in Figure. 1.

**Figure 1.**
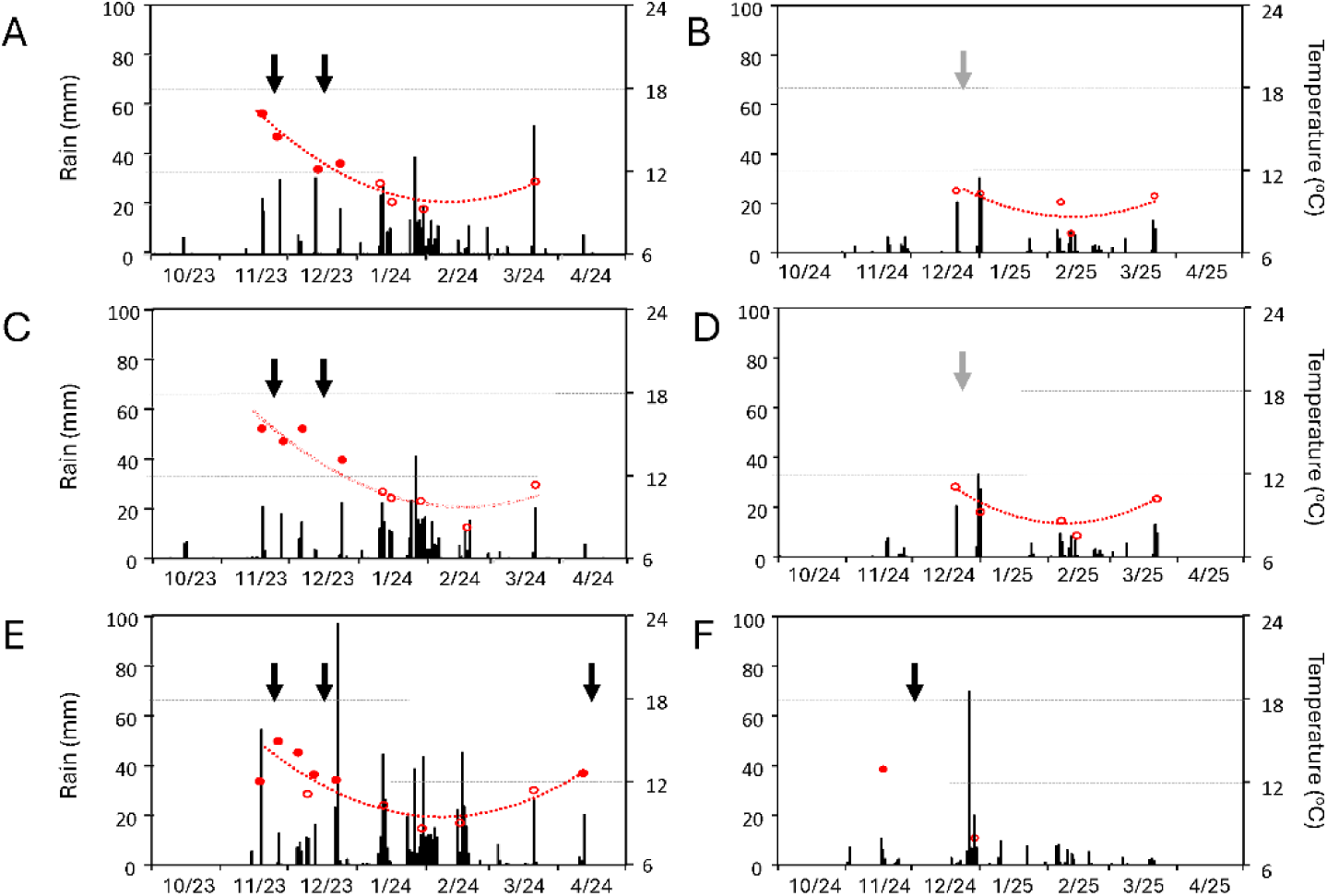
Weather data recorded at weather stations located at the vicinity of the experimental sited in the 2023/4 (A, C and E) and 2024/5 (B and D and F) growing seasons. **A** and **B** – experiments 1, 4 and 7; C and D experiments 2 and 5; E and F – experiments 3 and 6. Bars represent the daily quantity of rain. Circle symbol the average minimum temperatures in rain event with ≥ 15 mm of rain. Filled symbols: conditions fulfilled PedMan criteria for infection event. Empty symbols: conditions did not fulfill PedMan criteria for infection event. The red dashed lines represent the seasonal changes in minimum temperatures. Black arrows: timing of sprays applied according to PedMan recommendation; Gray arrows: timing of sprays applied in contrast to PedMan recommendation.

Our initial intention was to test the positive recommendations of PedMan (applying sprays in accordance with system’s recommendations), as outlined above. However, in the autumn months of the 2024/5 growing season, minimum temperatures during all rainy days in the sites of Kfar Ben Nun and Haruvit were below 12 °C. Accordingly, PedMan criteria for occurrence of infection event had not met and sprays should have not been applied. To evaluate the negative recommendation of the system (applying sprays in contrast to system’s recommendations) sprays were applied soon after one of the rain events with > 15 mm. These sprays were applied on 23 Dec 2024 (experiment 4) and 26 Dec 2024 (experiments 5 and 7) (Figure 1).

Experiments 1, 2, 4, 5 and 7 were laid out in randomized blocks and experiments 3 and 6 were laid out in completely randomized design. Each treatment was repeated four to five times, and each replication consisted of 1 tree. The fungicides used were ’Skipper’ (difenoconazole, EC 250 g/liter) produced and distributed in Israel by Tapazol Chemical Works, Ltd. (Beit Shemesh, Israel) and ’Nechostan’ (copper 190 g/liter, as 340 g/liter tribasic sulfate) produced by Nufarm and distributed in Israel by Adama- Agan (Ashdod, Israel). These fungicides are registered in Israel for peacock eye management and are commonly used by growers; they were sprayed in a tank-mix at concentrations of 0.06% and 0.3%, respectively. Fungicides were applied by means of a handgun at a volume of 15 liters water per tree.

#### Disease evaluation

Peacock eye disease affects tree health mostly by promoting leaf abscission (Graniti 1993). Incidence of leaf abscission (an indicator for plant health, in %) was visually estimated by two individuals (the same two in all assessments) at four canopy orientations of the experimental trees. Score of 0% implies that all leaves remained intact, and the trees seemed healthy with full, dense canopy; score of 50% implies that about half of the leaves had abscised and the trees’ canopy seemed transparent; score of 100% implies that all the leaves had abscised, the trees seemed as being completely defoliated (Ygzao et al. 2026c). Assessments in experiment 3 were performed in 2-to 3-week intervals starting in November and continuing until mid-July. The data were used to calculate the area under the leaf abscission progress curve (*AULAPC*) according to the following formula:

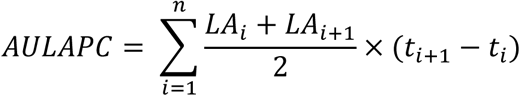

Where: *LAi* = incidence of leaf abscission (%) in time *i*; *ti* = assessment date (*t1* is the date of the first assessment and *tn* is the date of the last assessment); and *n* = number of assessments.

The relationship between the season-long assessment of leaf abscission (AULAPC) and the single assessment conducted in early May, corresponding to the peak of the second disease episode, was highly significant (*P* = 0.0005). The assessment conducted at the peak of the second disease episode explained 88.5% of the variation in season-long leaf abscission. Therefore, throughout this report, leaf abscission data collected at the peak of the second disease episode are presented. It should be noted that leaf abscission may result not only from peacock eye disease but also from other biotic factors, including fungal pathogens and insect pests, as well as abiotic stresses such as drought, waterlogging, salinity, nutrient imbalances, and temperature extremes. These factors can impair physiological functioning or damage leaf tissues, thereby inducing premature senescence and leaf abscission (Martin and Sibbett, 2005). Furthermore, because the assessors evaluated the entire tree rather than individual branches, it was not always possible to distinguish between leaf abscission occurring during the current season and that resulting from the previous season. Accordingly, within each experiment, differences in leaf abscission between fungicide-treated and untreated trees were attributed primarily to the effects of peacock eye disease. In each experiment, control efficacy (*CE*, %) was calculated by comparing the incidence of leaf abscission in the PedMan treatment (*LAₜ*) with that in the untreated control (*LAc*) using the following formula:

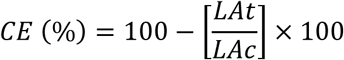

### Implementation of PedMan recommendations in commercial olive groves by growers

#### Observation procedures and treatments

During the 2025/26 growing season, PedMan recommendations were implemented by two olive growers in commercial groves located in two different production regions. The groves had not been sprayed against peacock eye disease in previous seasons, and the Souri trees grown in these groves were severely infected by the disease. The first grove was located in the Haruvit production area, in the vicinity of experiments 2 and 5. This area (hereafter referred to as Observation A) comprised eight olive grove subplots. In four of these subplots, the row orientation was north–south (subplots 31–34; central coordinates: 31.746329 N, 34.848459 E), while in the other four subplots the row orientation was east–west (subplots 41–44; central coordinates: 31.747569 N, 34.853523 E). The total area of the grove was 40 ha. In each subplot, the grower implemented the PedMan recommendation on a single row consisting of 20 trees and applied two spray treatments on 11 November and 14 December 2025 (Figure 2A). The second grove (hereafter referred to as Observation B) was located in Kfar Kish, at the site of experiments 3 and 6 (central coordinates: 32.668617 N, 35.465917 E). The total area of this grove was 1.2 ha. The grower implemented the PedMan recommendation across the entire plot and applied two spray treatments on 19 November and 23 December 2025 (Figure 2B). In both groves, the growers applied a mixture of Skipper and Nechostan (0.06% and 0.3%, respectively) using a Degania axial fan sprayer at a spray volume of 2,000 L/ha.

**Figure 2.**
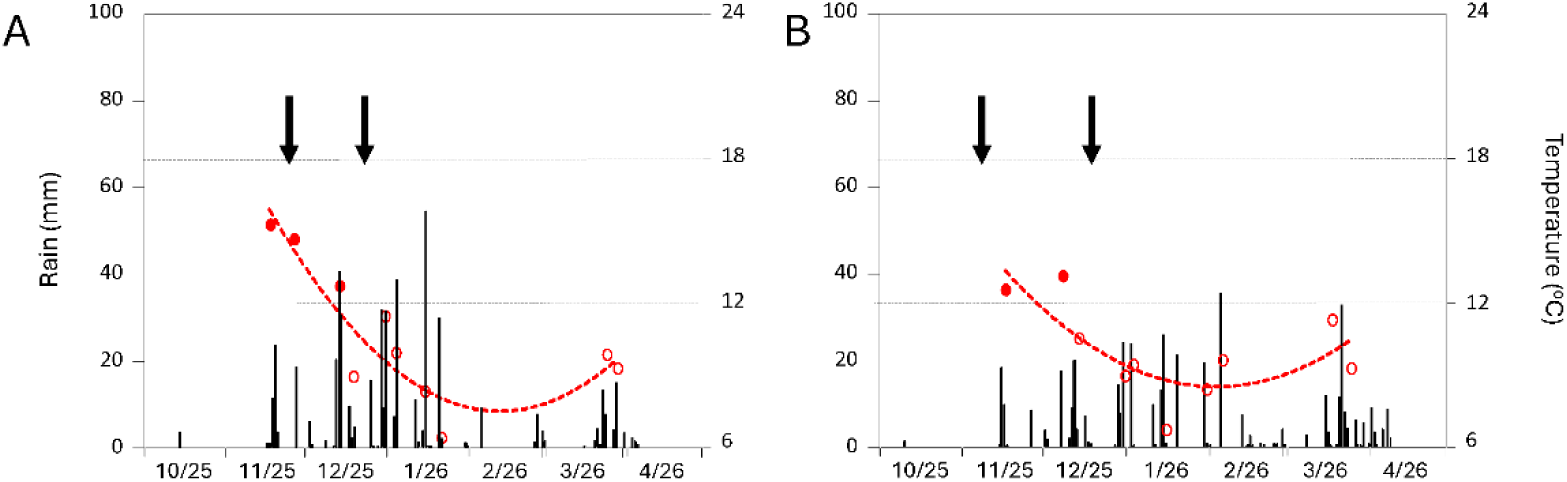
Weather data recorded at weather stations located at the vicinity of the observation sited in the 2025/6 growing season. **A** – observation A; **B** – observation B. Bars represent the daily quantity of rain. Circle symbols – the average minimum temperatures in rain event with ≥ 15 mm of rain. Filled symbols: conditions fulfilled PedMan criteria for infection event. Empty symbols: conditions did not fulfill PedMan criteria for infection event. The red dashed lines represent the seasonal changes in minimum temperatures. Black arrows: timing of sprays applied according to PedMan recommendation.

#### Evaluation of PedMan recommendations in the observations

The effects of the treatments on tree health were evaluated by assessing the incidence of leaf abscission (%) as described above. In observation A, leaf abscission was assessed on 10-20 trees in each of the eight subplots within the PedMan-sprayed rows. For comparison, similar assessments were conducted on cv. Souri trees located four rows away. These rows had not been sprayed against peacock eye disease and served as untreated controls. As noted above, the grower in observation B implemented the PedMan recommendations across the entire grove; therefore, no untreated control trees were available in this observation. Accordingly, trees included in experiment 6 were assessed; these trees had been subjected to a different spray history in previous growing seasons, as described below. In both observations assessments were performed in early May 2026 corresponding to the peak of the second disease episode (Ygzao et al. 2026c).

### Using PedMan for ascertaining various aspects related to peacock eye disease development

As mentioned above, PedMan predicts the occurrence of infection events was based on daily rain quantity (in mm) and the average minimum temperature (in °C) during the rainy days. Weather data for 11 weather stations located in the main olive production region in Israel were obtained for the 2020/1 to 2023/4 growing seasons (www.agrometeorology.co.il). The location of the weather stations is presented in Figure 3: Weather stations A and B are located in the Inner Plains, station C is located in Jezreel Valley, station D in Zevulun Valley, stations E and F in the Upper Galilee Mountains, station E in the Southern Sea Shore, station H in the Lower Galilee Mountains, station I in Golan Heights, station J in Beit-She’an Valley and station K in the Northern Negev region. According to Köppen-Geiger climate classification (Peel et al. 2007), the climate in sites A to I is classified as Csa – hot summer Mediterranean climate; the climate in site J is classified as BSh – hot semi-arid and the climate in site K is classified as BWh – hot desert climate (https://ims.gov.il).

**Figure 3.**
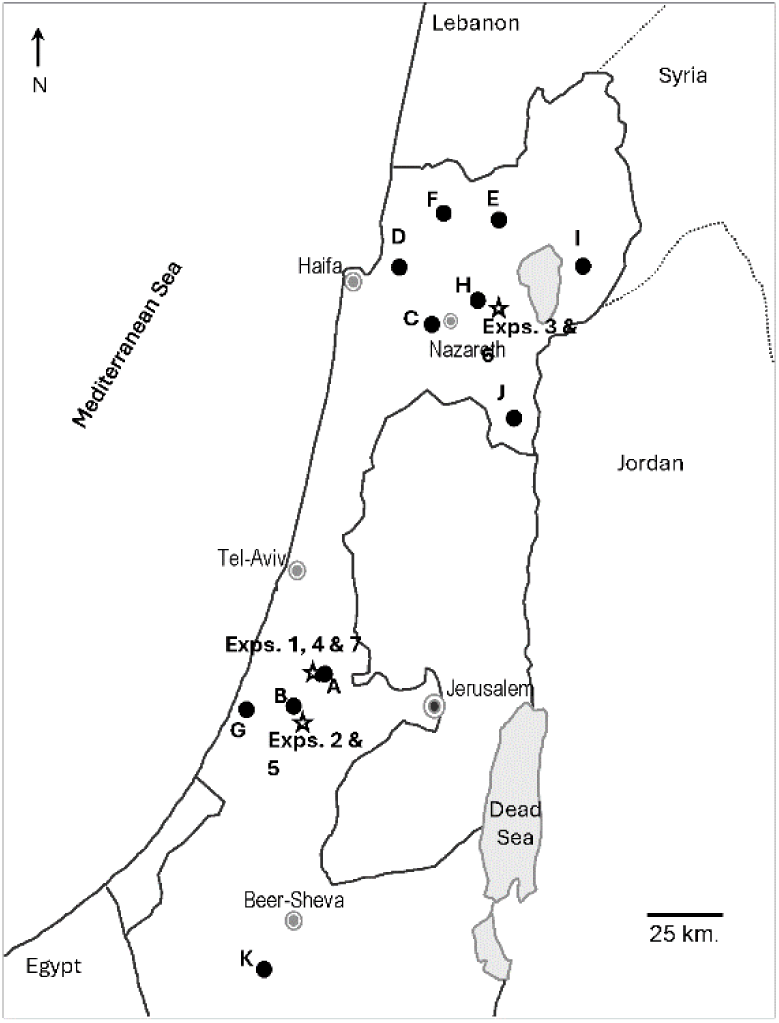
The location of weather stations (marked in filled circles and named A to K) from which weather data were obtained for running the PedMan system. Weather stations A and B are located in the Inner Plains, station C is located in Jezreel Valley, station D in Zevulun Valley, stations E and F in the Upper Galilee Mountains, station E in the Southern Sea Shore, station H in the Lower Galilee Mountains, station I in Golan Heights, station J in Beit-She’an Valley and station K in the Northern Negev region. The sites of experiments 1 to 7 are presented by open stars.

The weather data obtained from the weather stations were used to perform simulation runs in which PedMan identified the occurrence and timing of infection events. The results were used for ascertaining various aspects related to peacock eye disease development, as follows: (i) Previous results suggested that infection events in Israel occur during the autumn and spring months (Ygzao et al. 2026a; b) but it is not known when and how often infection events occur. PedMan predictions were used to determine the monthly distribution of infection events and the partitioning between the autumn and spring months. (ii) Olive groves are scattered across Israel in regions with dissimilar weather conditions. Although there is a general information on the intensity of peacock eye disease in the different regions, this knowledge is inferred from laboratory or greenhouse studies extrapolation and not on real-world commercial experiments under different climatic conditions. The number of infection events in the autumn months of each season may serve as an indicator for epidemic potential in that growing season. PedMan predictions were used to categorize the olive production areas in Israel into groups according to the potential severity of disease epidemics. Furthermore, it enabled to estimate the variation in the potential severity of the disease among seasons. (iii) The number of infection events in the spring months of each season may serve as an indicator for the survival potential of the pathogen between growing seasons. PedMan predictions were used to categorize the olive production areas into groups according to the survival potential of the pathogen. Furthermore, it enabled to estimate the variation in the survival potential of the pathogen among spring seasons. (iv) The likelihood for severe epidemic is the highest when both, the survival potential of the pathogen between growing seasons and the epidemic potential in the growing season, are high. PedMan predictions were used to categorize the olive production areas into groups based on the likelihood for occurrence of severe epidemics. (v) According to PedMan criteria, the shortest interval between sprays is 14 days. Accordingly, the seasonal number of sprays scheduled by the system could be equal, or smaller, to the number of predicted infection alerts. PedMan predictions were used to estimate the average and the maximal number of sprays that could have been scheduled by the system in the autumn and in the spring months.

## Data analyses

As indicated above, the experiments in Kfar Kish (experiment 3 and 6) were carried out in the same site where another experiment was performed in 2022/3. Accordingly, in 2024/5 there were trees that were diversly sprayed in each of the growing season. There were six treatments in total (each with 4- 5 replicates), as follows: **1**. Trees that were not sprayed with fungicides in none of the growing seasons; **2-4**. Trees that were sprayed only in one of the growing seasons, 2022/3, 2023/4 or 2024/5; **5**. Trees that were not sprayed in 2022/3 but were sprayed in the 2023/4 and 2024/5 growing seasons; and **6**. Trees were sprayed with fungicides in all three growing seasons. To quantify the contribution of spraying in each of the growing seasons on the incidence of leaf abscission at the peak of the second disease episode of 2024/5, we employed multiple regression with dummy variables, a general statistical procedure which encompass Analysis of Variance (Netter et al. 1985). This technique is useful in identifying quantitative effects of variables having distinct levels. We assumed that the effect of spraying in each growing season was additive. Accordingly, we employed an additive model to quantify the effects of spraying on tree health. The following regression model was used:

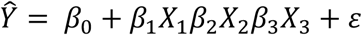

In which: *Ŷ* is the dependent variable denoted to the incidence of leaf abscission (in %) at the peak of the second disease episode of 2024/5; *β*0 to *β*3 are the regression coefficients; X1 to X3 are the independent variables denoted to spraying in the 2022/3 (X1), 2023/4 (X2) or 2024/5 (X3) growing seasons; and ε is the error term. In multiple regression with dummy variables, the actual value of each independent variable is either 0 or 1. A value of 0 was assigned for treatments where sprays were not applied in a certain growing season and value of 1 for treatments where sprays were applied. The timing of spraying in the 2022/3 growing season was presented elsewhere (Ygzao et al. 2026b); sprays in 2023/4 and 2024/5 growing seasons were timed according to PedMan recommendations (Figure 1E and F).

Throughout the study, we used JMP 16 Pro software to conduct statistical analyses. The criteria were based on optimizing the Akaike Information Criterion (AIC), Bayesian Information Criterion (BIC), or mean Squared Error (MSE), depending on the context of the model. Figures and visual representations were created using Microsoft Excel.

## Results

### Evaluating the performance of PedMan in grove experiments under natural conditions

The positive recommendations generated by PedMan (*i.e*., fungicide applications made according to the system’s recommendations) were implemented for the first time in four experiments (Experiments 1, 2, and 3 during the 2023/24 growing season, and Experiment 6 during the 2024/25 growing season). Applying fungicides according to PedMan recommendations had only a negligible effect on leaf abscission during the first disease episode (data not shown). In contrast, during the second disease episode, the incidence of leaf abscission was markedly lower in PedMan-treated trees than in the untreated control (Figure 4a and B; Table 1). Across the four experiments, the incidence of leaf abscission in PedMan-treated trees was significantly reduced by 60.0 ± 5.3% (mean ± SE) relative to the untreated control.

**Figure 4.**
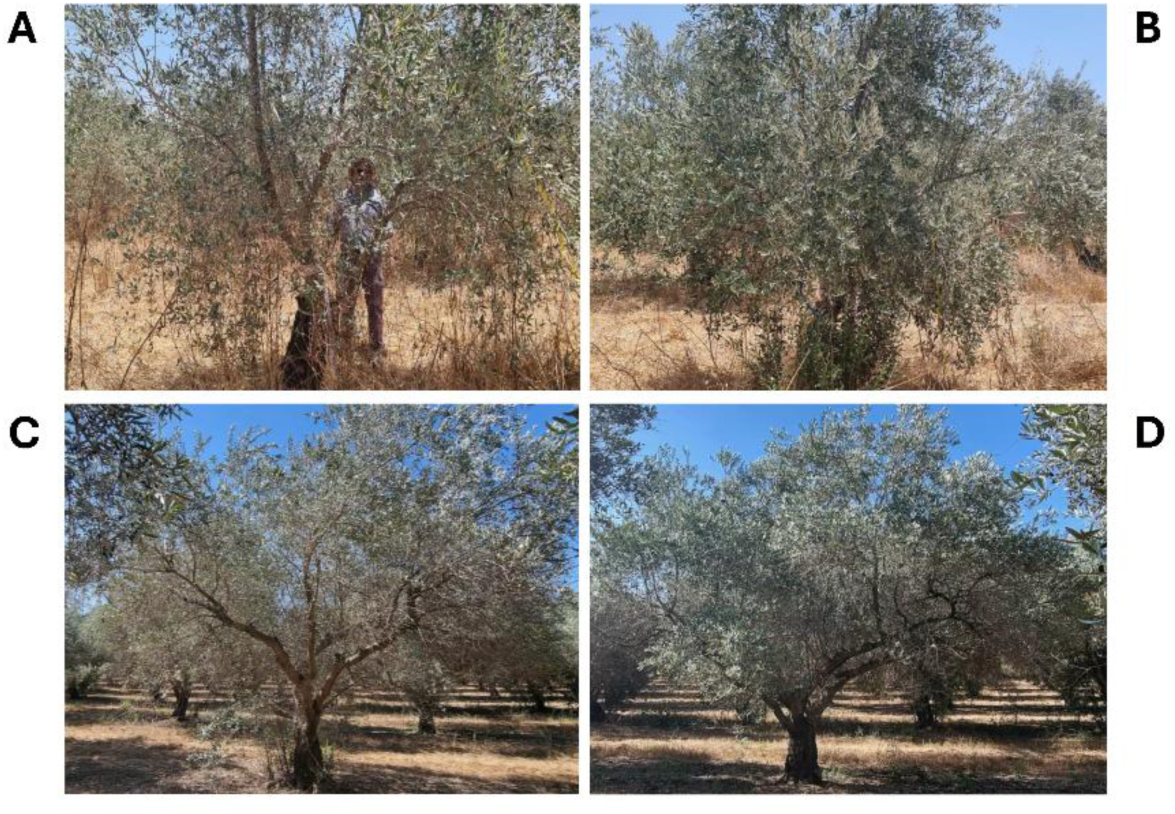
A and **B**: Photos taken on late April at the peak of the second disease episode of the 2023/4 season in experiment 1. DE is standing behind the trees in both photos to demonstrate the transparency of the trees as result of leaf abscission. **C** and **D**: Photos taken on late April at the peak of the second disease episode of the 2025/6 season in observation A. **A** and **C**: Untreated control; **B** and **D**: The PedMan treatment.

**Table 1.** The incidence of leaf abscission in the experiments where sprays were applied in accordance to PedMan advice to evaluate the positive recommendation of the system^y^

| Treatment in: |  | Experiment number and growing season: |  |  |  |
|---|---|---|---|---|---|
| Previous season | Current season | Exp. 1 (23/4) | Exp. 2 (23/4) | Exp. 3 (23/4) | Exp. 6 (24/5) |
| Control | Control | 29.0 a<sup>z</sup> | 37.5 a | 33.8 a | 27.4 a |
| Control | PedMan | 6.9 b | 14.9 b | 18.1 b | 11.8 b |
| PedMan | Control | - | - | - | 21.7 a |
| PedMan | PedMan | - | - | - | 7.5 b |

<sup>y</sup>The incidence of leaf abscission (%) was evaluated at the peak of the second disease episode.
<sup>z</sup>In each column, numbers followed by the same letter do not differ significantly (at *P* ≤ 0.05) as determined by HSD test.

In Experiment 6 (2024/25), PedMan recommendations were followed for two consecutive growing seasons, resulting in an even greater reduction in leaf abscission (72.6%). In contrast, when PedMan recommendations had been implemented in the previous growing season but not in the current one, the incidence of leaf abscission was only slightly lower (20.8%) than in the untreated control, and this reduction was not statistically significant (Table 1).

Minimum temperatures during the rainy days in the sites of experiments 4, 5 and 7 in the autumn months of 2024/5 growing season, were below 12 °C (Figure 1B and D). Accordingly, PedMan criteria for occurrence of infection event had not met and sprays should have not been applied. To evaluate the negative recommendation of the system (applying sprays in contrast to system’s recommendations) sprays were applied soon after the occurrence of ≥ 15 mm of rain. In all experiments, differences in the incidence of leaf abscission between the assessments performed during January and February, at the peak of the first disease episode and the assessments performed during April and May, at the peak of the second disease episode were minute, indicating that leaf abscission was not intensified in the experimental sites. Furthermore, in all experiments the differences in the incidence of leaf abscission between the PedMan treated and the untreated control trees were insignificant, indicating that the sprays applied in contrast to system’s recommendations did not improve tree health (Table 2).

**Table 2.** The incidence of leaf abscission in the experiments where sprays were applied in in contrast to PedMan advice to evaluate the negative recommendation of the system^y^

| Treatment in: |  | Exp. 4 (24/5) |  | Exp. 5 (24/5) |  | Exp. 7 (24/5) |  |
|---|---|---|---|---|---|---|---|
| Previous season | Current season | 1<sup>st</sup> disease episode | 2<sup>nd</sup> disease episode | 1<sup>st</sup> disease episode | 2<sup>nd</sup> disease episode | 1<sup>st</sup> disease episode | 2<sup>nd</sup> disease episode |
| Control | Control | 25.5 a<sup>z</sup> | 26.4 a | 25.2 a | 25.3 a | 38.9 a | 36.7 a |
| Control | PedMan | - | - | - | - | 37.0 a | 37.0 a |
| PedMan | PedMan | 16.1 a | 15.9 a | 19.5 a | 19.4 a | - | - |

<sup>y</sup>The incidence of leaf abscission was evaluated during January and February or at April and May, at the peak of the first or the second disease episodes, respectively.
<sup>z</sup>In each column, numbers followed by the same letter do not differ significantly (at *P* ≤ 0.05) as determined by HSD test.

Multiple regression analysis with dummy variables was used to estimate the impact of spraying in each of the previous three growing seasons on tree health in experiment 6, at the peak of the second disease episode of 2024/5. Spraying according to PedMan in the 2023/4 and 2024/5 growing seasons reduced the incidence of leaf abscission in 2024/5 significantly; spraying in 2022/3 also reduced the incidence of leaf abscission but this effect was insignificant (Table 3). Results of the regression analysis enabled to estimate the outcome of spraying at different scenarios on the incidence of leaf abscission. Spraying in only one of the growing seasons reduced the incidence of leaf abscission by 13.3-52.2%; The largest reduction was obtained when the sprays were applied in the current growing season. Spraying in two growing seasons reduced the incidence of leaf abscission by 33.3-72.2%; The largest reduction was obtained when the sprays were applied in the current and previous growing seasons. Spraying in the previous three growing seasons reduced the incidence of leaf abscission by 85.5% (Table 4).

**Table 3.**
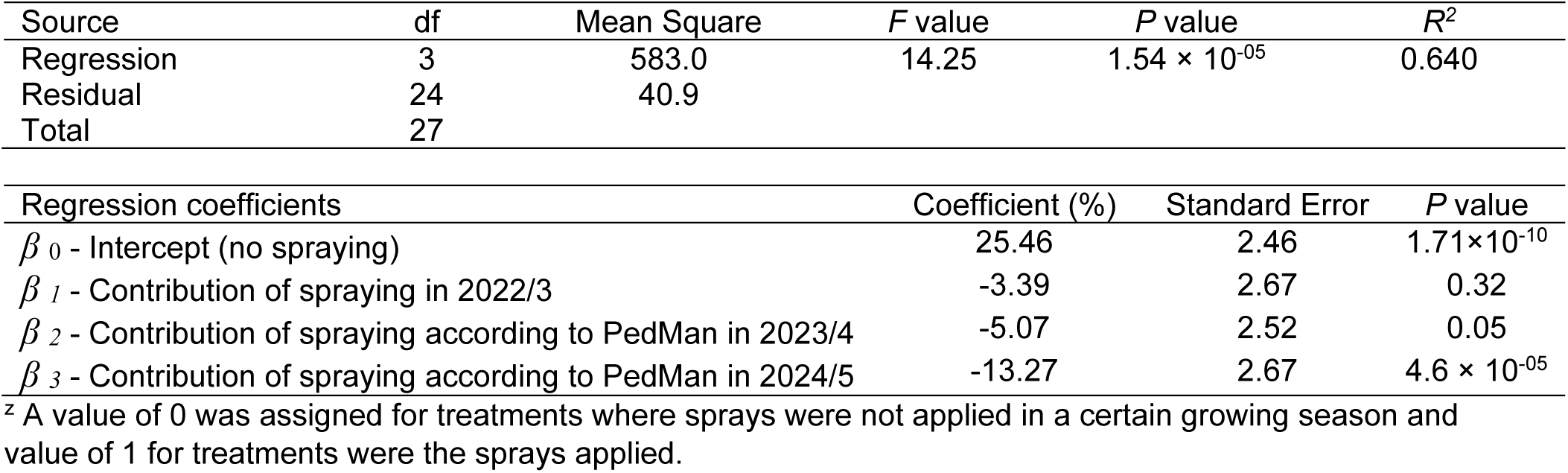
Analysis of variance of the multiple regression analysis with dummy variables performed to estimate the effects of spraying in three growing seasons on the incidence leaf abscission in experiment 3, at the peak of the second disease episode of 2024/5^z^

| Source | df | Mean Square | *F* value | *P* value | *R<sup>2</sup>* |
|---|---|---|---|---|---|
| Regression | 3 | 583.0 | 14.25 | 1.54 × 10<sup>-05</sup> | 0.640 |
| Residual | 24 | 40.9 |  |  |  |
| Total | 27 |  |  |  |  |
| Regression coefficients |  |  | Coefficient (%) | Standard Error | *P* value |
| *β*<sub>*0*</sub> - Intercept (no spraying) |  |  | 25.46 | 2.46 | 1.71×10<sup>-10</sup> |
| *β*<sub>*1*</sub> - Contribution of spraying in 2022/3 |  |  | -3.39 | 2.67 | 0.32 |
| *β*<sub>*2*</sub> - Contribution of spraying according to PedMan in 2023/4 |  |  | -5.07 | 2.52 | 0.05 |
| *β*<sub>*3*</sub> - Contribution of spraying according to PedMan in 2024/5 |  |  | -13.27 | 2.67 | 4.6 × 10<sup>-05</sup> |

<sup>z</sup> A value of 0 was assigned for treatments where sprays were not applied in a certain growing season and value of 1 for treatments were the sprays applied.

**Table 4.**
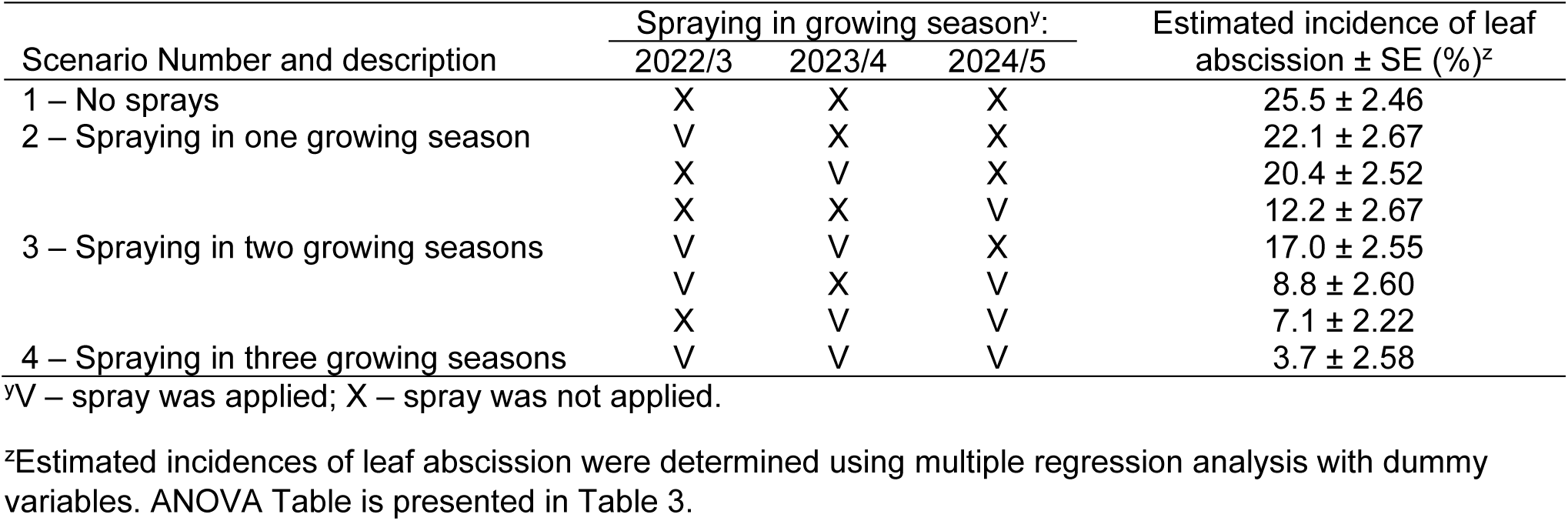
Effects of various scenarios of spraying in three growing seasons (2022/3 to 2024/5) on the incidence of leaf abscission in experiment 3, at the peak of the second disease episode of 2024/5

| Scenario Number and description | Spraying in growing season<sup>y</sup>: |  |  | Estimated incidence of leaf abscission ± SE (%)<sup>z</sup> |
|---|---|---|---|---|
|  | 2022/3 | 2023/4 | 2024/5 |  |
| 1 – No sprays | X | X | X | 25.5 ± 2.46 |
| 2 – Spraying in one growing season | V | X | X | 22.1 ± 2.67 |
|  | X | V | X | 20.4 ± 2.52 |
|  | X | X | V | 12.2 ± 2.67 |
| 3 – Spraying in two growing seasons | V | V | X | 17.0 ± 2.55 |
|  | V | X | V | 8.8 ± 2.60 |
|  | X | V | V | 7.1 ± 2.22 |
| 4 – Spraying in three growing seasons | V | V | V | 3.7 ± 2.58 |

<sup>y</sup>V – spray was applied; X – spray was not applied.
<sup>z</sup>Estimated incidences of leaf abscission were determined using multiple regression analysis with dummy variables. ANOVA Table is presented in Table 3.

### Implementation of PedMan recommendations in commercial olive groves by growers

PedMan recommendations were implemented by olive growers in their commercial groves in two production areas. In both cases, implementing PedMan recommendations significantly decreased leaf abscission and improved tree health (Figure 4C and D; Table 5).

**Table 5.**
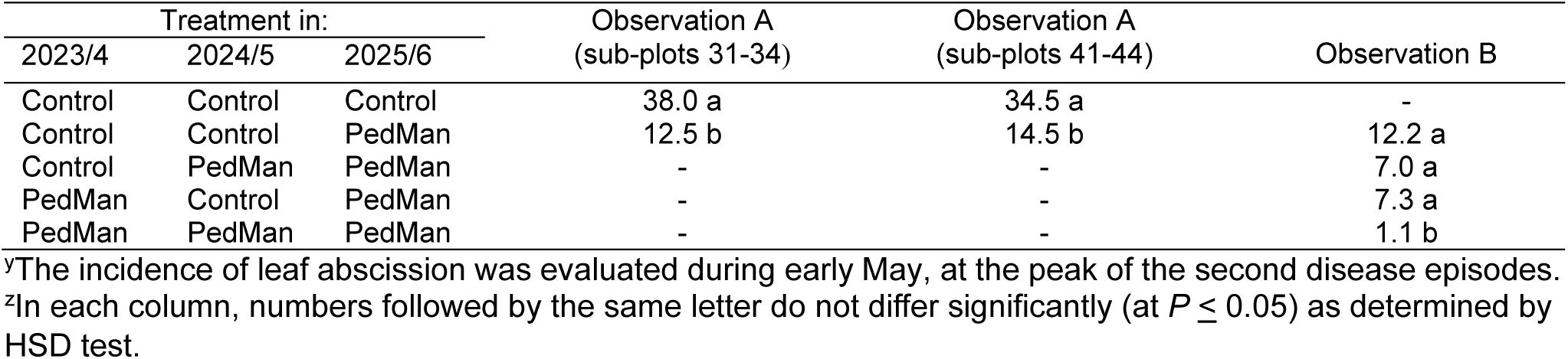
The incidence of leaf abscission in in commercial olive groves where sprays were applied in the 2025/6 growing season in accordance to PedMan advice to evaluate the positive recommendation of the system^y^

| Treatment in: |  |  | Observation A<br>(sub-plots 31-34) | Observation A<br>(sub-plots 41-44) | Observation B |
|---|---|---|---|---|---|
| 2023/4 | 2024/5 | 2025/6 |  |  |  |
| Control | Control | Control | 38.0 a | 34.5 a | - |
| Control | Control | PedMan | 12.5 b | 14.5 b | 12.2 a |
| Control | PedMan | PedMan | - | - | 7.0 a |
| PedMan | Control | PedMan | - | - | 7.3 a |
| PedMan | PedMan | PedMan | - | - | 1.1 b |

<sup>y</sup>The incidence of leaf abscission was evaluated during early May, at the peak of the second disease episodes.
<sup>z</sup>In each column, numbers followed by the same letter do not differ significantly (at *P* ≤ 0.05) as determined by HSD test.

### Using PedMan for ascertaining various aspects related to peacock eye disease development

After evaluating PedMan recommendations under commercial field conditions, the system was used in simulation runs for ascertaining various aspects related to peacock eye disease development. First, the timing of infection alerts was determined. Most of the infection alerts (77.8%) occurred in the autumn and early winter months (September to December) with November being the most common month. Few infection alerts occurred in the mid-winter months (January and February) and only 14.8% of the seasonal alerts occurred in the spring months, most of them in March and April (Figure 5).

**Figure 5.**
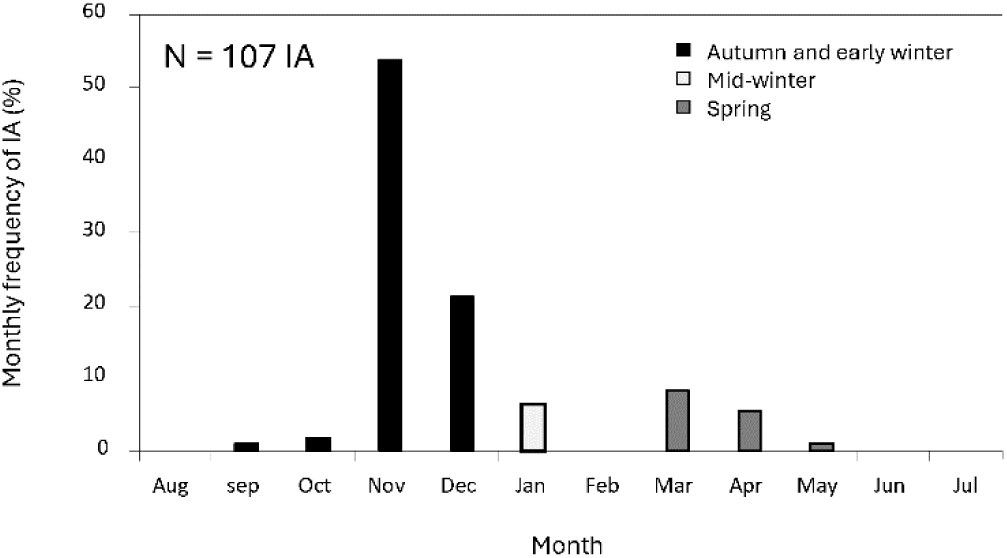
The monthly distribution of infection alerts (IA) that were predicted by PedMan using for weather data recorded in 11 stations located in the main olive production areas of Israel in the growing seasons of 2020/1 to 2023/4.

The number of infection alerts in the autumn and spring months varied markedly between the sites and in each site, between the growing seasons (Figure 6). Analysis of this variation enabled to categorize the olive production areas into three groups. The number of infection alerts in the Inner Plains and Jezreel Valley (sites A, B and C; Figure 3) was the largest, with three or more alerts on the average in a growing season. The number of infection alerts in Zevulun Valley, Upper Galilee Mountains and the southern Sea Shore (sites D, E-F, and G; respectively) was modest with 1.5 to 3 alerts on the average. In these sites, there was at least one infection alert in each growing season (Figure 6A). The number of infection alerts in the Lower Galilee Mountains, Golan Heights, Beit-She’an Valley and the Northern Negev region (sites H, I, J and K, respectively) was the least with less than 1.5 infection alerts on the average. In these sites, in 9 out of the 16 cases examined (56.2%) there were no alerts at all in the autumn months (Figure 6A). As expected, there was a considerable variation in the number of infection alerts among growing seasons. Whereas the number of alerts in the autumn months of the 2020/1 growing season was 3.2, on the average, in the 2021/2 growing season there were only 0.8 events (Figure 6B).

**Figure 6.**
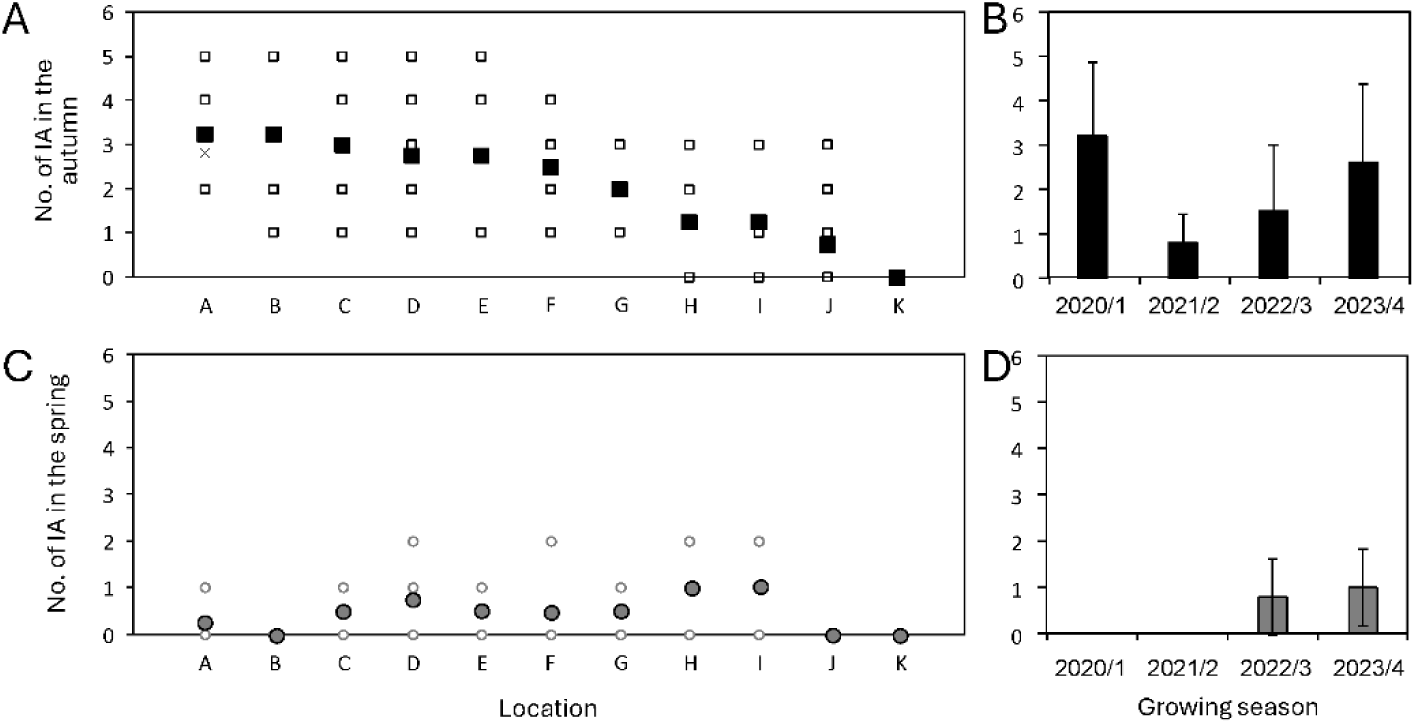
The number of infection alerts (IA) predicted by PedMan using weather data recorded in 11 stations located in the main olive production areas of Israel in the growing seasons of 2020/1 to 2023/4 for the autumn (**A** and **B**) and spring months (**C** and **D**). Open symbols in graphs **A** and **C** denote to the yearly number and filled symbols to the mean number of infection alerts in each location. Vertical bars in graphs **B** and **D** denote to the seasonal SD. The location of the weather stations is presented in Figure 3.

The number of infection alerts in the spring months in sites D, H and I (Figure 6C) was the largest, with 0.75 or more alerts, on the average. The number of infection alerts sites A, C and E-G was modest with 0.25 to 0.5 alerts on the average. No infection alerts were issued in sites B, J and K (Figure 6C). It should be noted that in the 2020/1 and 2021/2 growing seasons there were no infection alerts in the spring months in all sites (Figure 6D).

The likelihood for severe epidemic is the highest when both, the survival potential of the pathogen between growing seasons and the epidemic potential in the growing season, are high. In 6 out of the 33 cases examined (18.2%) there were infection alerts in the spring months of one growing season and in the autumn months of the succeeding growing season. These situations occurred in 10 sites. Conversely, the likelihood for mild epidemic is the highest when both, the survival potential of the pathogen between growing seasons and the epidemic potential in the growing season, are low. In 8 out of the 33 cases examined (24.2%) there were no infection alerts in the spring months of one growing season and in the autumn months of the succeeding growing season. This situation occurred only in sites H, I, J and K (Figure 6). In the rest of the cases (19 cases; 57.6%) there were no infection alerts in the spring months of one growing season but there were some (one or more) infection alerts in the autumn months of the succeeding growing season. In these cases, it is expected that epidemics will develop, but their intensity will be modest. This situation occurred in 10 of the sites.

The relationship between the number of infection alerts issued by PedMan and the number of sprays recommended by the system was highly significant (*R^2^* = 0.968; *P* < 0.0001). The number of scheduled sprays varied between growing seasons and sites (between 0 and 4; results not shown) but on the average, the number of sprays scheduled in the autumn months did not exceed 2 and the number of sprays scheduled in the spring months did not exceed 1 (Figure 7).

**Figure 7.**
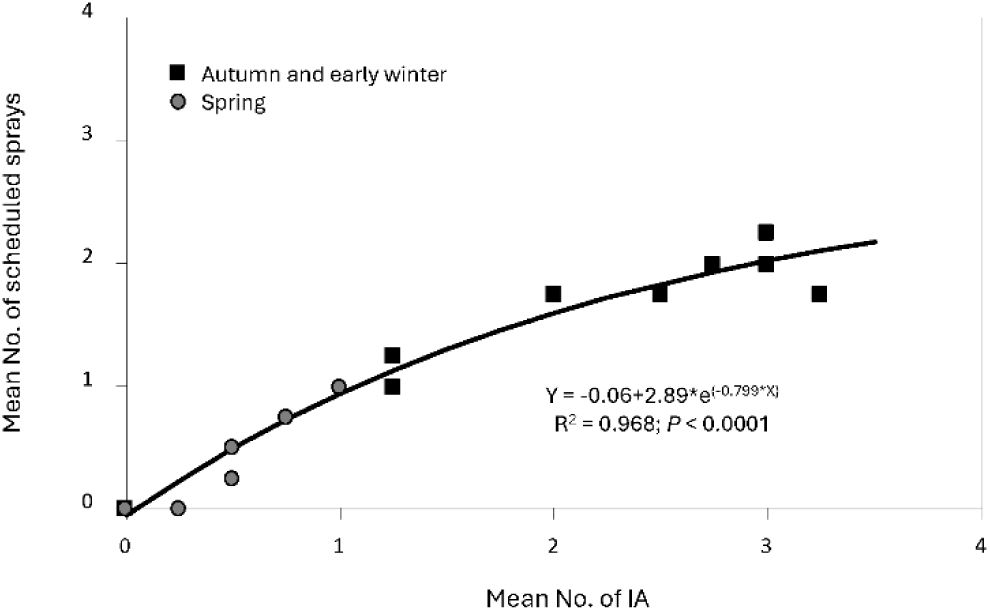
The relationship between the mean number of infection alerts (IA) predicted by PedMan and the mean number of sprays scheduled by the system. Weather data were recorded in 11 stations located in the main olive production areas of Israel in the growing seasons of 2020/1 to 2023/4.

## Discussion

The performance of forecast models and DSSs predictions must be evaluated under field conditions to determine the occurrence of true (or false) positive and true (or false) negative predictions (Thomidis et al. 2021). In this study, we report the results of four experiments aimed at evaluating the true positive recommendations of PedMan. In all experiments, the application of fungicides according to the system significantly improved tree health, confirming that PedMan recommendations to spray were accurate (Table 1). In three experiments, PedMan predicted that infection events had not occurred. In these experiments, fungicide applications were nevertheless performed to assess the negative recommendations of the system. In all cases, applying fungicides contrary to the system’s recommendation had no effect on tree health, confirming that PedMan recommendations not to spray were also accurate (Table 2). In the following growing season, the system was used by growers in their commercial groves and the results were satisfactory (Table 5). These results indicate that the PedMan successfully identifies conditions under which spraying is beneficial, supporting its utility as a decision-support tool.

As indicated above, a considerable number of forecast models have been developed to predict *V. oleaginea* infections and peacock eye disease development. However, only a limited subset of these models has undergone rigorous validation under independent field conditions. In most cases, model performance has been assessed using the same or closely related datasets employed during model development (*i.e.,* verification), raising concerns regarding their generalizability and robustness across diverse agroecological contexts. Independent field evaluation of these models (*i.e.,* validation) remains scarce, with studies such as Thomidis et al. (2021) representing one of the few efforts that systematically examined forecast model predictions using independent data originating from commercial groves. This gap highlights the need for additional independent, multi-site validation studies to ensure the reliability and practical applicability of forecast models and DSSs. Furthermore, the primary motivation underlying the development of most forecast models and DSSs has been to improve the timing of fungicide applications (Buonaurio et al. 2023; Rhimini et al. 2020; Romero et al. 2018; Roubal et al. 2013; Thomidis et al. 2021; Viruega and Trapero 1999). As indicated above, studies reporting grove experiments in which forecast model or DSS predictions were used to guide spray timing have not been published. Accordingly, it appears that PedMan is the first DSS to have been thoroughly validated for timing fungicide applications against peacock eye disease.

Two of our experiments were conducted in the same groves over two or three consecutive growing seasons. Analysis of the results enabled estimation of the cumulative contribution of fungicide applications applied over one, two, or three successive growing seasons to improvements in tree health. The results showed that the contribution of multi-season spraying to reducing the incidence of leaf abscission increased progressively: from 60.8%, on the average, when sprays were applied only in the current growing season, to 72.6% when sprays were applied in both the current and the preceding growing seasons, and to 85.5% when sprays were applied over three consecutive growing seasons (Tables 1 and 4). Assuming that fungicide applications are continued in subsequent seasons and that their cumulative effect follows a similar logistic trend, it is expected that after five consecutive years of spraying, the incidence of leaf abscission would become negligible and infected trees would fully recover. Conidia of *V. oleaginea* are disseminated during rain events by raindrops; therefore, their dispersal is typically limited to the distance traveled by raindrops. Consequently, new infections generally originate within the same tree canopy or from adjacent trees located only a few meters away (Issa et al. 2019; Viruega et al. 2013; Wilson and Miller 1949). This localized dispersal suggests that the pathogen could potentially be eradicated from infected olive groves if fungicide applications are properly timed over five consecutive growing seasons. However, when trees were treated in one growing season but not in the following season, the incidence of leaf abscission increased by 60.2– 78.2% (Tables 1 and 4). These findings highlight the polyetic nature of the pathogen and emphasize the necessity for continuous, multi-season disease suppression.

Conclusions derived from our grove experiments were corroborated and further expanded through analysis of the simulation runs conducted in this study. It is well established that *V. oleaginea* infections occur under conditions of frequent rainfall, prolonged leaf wetness, and mild temperatures. In both the grove experiments and the simulation runs, most infection events occurred during the autumn and early winter months. Some infection events were observed in spring, whereas very few, if any, occurred during the winter and summer months (Figures 1, 2 and 5). These findings are consistent with existing recommendations to apply fungicides during the autumn and spring (Almadi et al. 2024; Obanor et al. 2005; Roca et al. 2010; Trapero and Blanco 2010; Viruega et al. 2011). Our results help identify the key factors that promote or constrain infection events under the Mediterranean climate prevailing in Israel. In autumn and early winter, rainfall occurs while temperatures remain above the lower threshold for conidial germination (Figures 1 and 2; Ygzao et al. 2026a). Accordingly, rainfall is the primary factor governing the occurrence of infection events during this period. During winter, although rainfall and leaf wetness are abundant, temperatures during rainy periods are often too low to permit conidial germination and infection. Thus, low temperature is the principal factor limiting infection events in winter. In spring, rainfall occurs intermittently, but temperatures during rainy periods are often still below the threshold required for infection. However, when temperatures are sufficiently high, infection events may occur (Figure 1). Therefore, temperature is the main factor governing infection occurrence in spring. In contrast, summers in Mediterranean climates are typically hot and dry: rainfall is absent, leaf wetness is minimal, and temperatures exceed the upper threshold for conidial germination (Ygzao et al. 2026a). Consequently, the absence of rainfall, limited leaf wetness, and high temperatures collectively prevent infection events during summer. Considering this analysis, in olive production areas with different climatic conditions, (for example, Csb - warm summer Mediterranean climate), or under future climate change scenarios (e.g., increased winter temperatures) (Fraga et al. 2021), the limiting factors described above may no longer apply, and infection events could occur during different periods of the year.

As indicated above, new infections generally originate within the same tree canopy or from adjacent trees located only a few meters away (Issa et al. 2019; Viruega et al. 2013; Wilson and Miller 1949). This localized dispersal implies that each olive grove functions as a distinct epidemiological unit with respect to peacock eye disease. Accordingly, the multi-season development of the disease depends on the pathogen’s ability to survive within the same grove. *V. oleaginea* survives the summer on infected leaves that remain attached to the tree; it does not persist on fallen leaves (Buonaurio et al. 2023; González-Domínguez et al. 2017; Viruega et al. 2013). These over-summering infected leaves serve as the primary source of inoculum in the following autumn (Ygzao et al. 2026c). Their abundance depends on the occurrence of infection events during the autumn and spring of the previous growing season (Ygzao et al. 2026c). Although there was considerable variation in the occurrence and intensity of infection events among growing seasons, such events were recorded in all olive production areas in Israel (Figures 1, 2 and 6). The only exception was site K, characterized by a BWh (hot desert) climate, where the scarcity of rainfall, low humidity, and high temperatures preclude the occurrence of infection events (Figure 6). These findings indicate that the pathogen can develop and persisting in most olive- growing regions of Israel. However, the actual severity of the disease is strongly influenced by the specific weather conditions prevailing in a given growing season. These conclusions are consistent with observations regarding the prevalence of the disease in olive groves across Israel (R. Birger and M. Yunis, *personal communication*).

In some countries, it has been recommended to adjust the number and timing of fungicide applications according to the conduciveness of prevailing weather conditions to peacock eye disease. It is recommended to apply two copper-based sprays in areas with high disease potential; the first in autumn, prior to the onset of the rainy season, and the second in spring. In areas with moderate disease potential, it is recommended to spray only in the autumn, whereas no sprays are recommended in areas with low disease potential (Almadi et al. 2024; Obanor et al. 2005; Roca et al. 2010; Trapero and Blanco 2010; Viruega et al. 2011). However, results from our grove experiments (Tables 1, 2 and 5) and analyses of the simulation runs (Figure 6) indicate that the conduciveness of weather conditions to peacock eye disease fluctuates markedly across growing seasons in most production areas. In some growing seasons, weather conditions promoted numerous infection events; in others, conditions were less favorable; and in certain growing seasons, conditions were insufficient for infection events to occur at all. A similar pattern was even observed at site J, characterized by a BSh (hot semi-arid) climate (Figure 6). Multi-season fluctuations in disease severity were reported by other authors in different countries (for example, Romero et al. 2018; Roubal et al. 2013; Viruega and Trapero 1999). Our findings suggest that in most olive-growing regions in Israel, it is not possible to classify a priori the conduciveness of weather conditions to the pathogen or to prescribe a fixed number of fungicide applications for a given growing season. Moreover, because the timing of infection events may vary widely within growing seasons - ranging from September to December for the autumn period and from March to May for the spring period (Figures 1, 2 and 5) and because fungicide efficacy is limited to approximately 14 days (Ygzao et al. 2026b), it is not feasible to define the optimal spray timing in advance. This conclusion underscores the need for forecast models or DSSs to determine the precise timing of fungicide applications in response to prevailing weather conditions. As demonstrated in the present study, PedMan successfully fulfilled this role.

The number of infection alerts issued by PedMan ranged from 0 to 5 during the autumn period of infection events and from 0 to 2 during the spring period of infection events (Figures 1, 2 and 6). Because the minimum interval between two consecutive sprays scheduled by PedMan is 14 days, the number of fungicide applications per season is expected to be equal to or lower than the number of infection events. Accordingly, the number of sprays scheduled by the system in our grove experiments and observations (Figures 1 and 2) and simulation runs (Figure 7) ranged from 0 to 3 per growing season, corresponding to the number of sprays recommended in other studies (Almadi et al. 2024; Belfiore et al. 2014; Buonaurio et al. 2023; Obanor et al. 2008; Roca et al. 2010). These findings indicate that although the use of PedMan enabled to improve the timing of fungicide application, it does not increase the number of fungicide applications compared with currently recommended spray programs in other countries.

Data originating from our laboratory and grove experiments conducted during the 2021/2 to 2022/3 growing seasons (Ygzao 2026a, b, c) provided the foundational knowledge required for the development of PedMan. The system was initially verified by its developers using the same data on which it was constructed (Ezra et al. 2026; Gilat et al. 2026). Subsequently, the system’s predictions were validated in seven grove experiments carried out during the 2023/4 and 2024/5 growing seasons. The results, presented in the current study, indicate that PedMan recommendations are accurate in highly susceptible cultivar. In the following growing season (2025/6), PedMan was implemented commercially by olive growers in two distinct olive production regions, allowing them to achieve effective disease suppression. It should be noted that these growers had not applied fungicides against peacock eye disease in previous growing seasons because, based on their past experience, such treatments were ineffective. In a parallel study (Yotam et al., 2026) we upgraded the system and evaluated its performance in additional cultivars. In our opinion, PedMan could also be used in other olive productions areas with a Csb climate (Fraga et al. 2021; Torres et al. 2017) in the Mediterranean countries and possibly also in other parts of the world. However, prior to its implementation in other regions, PedMan predictions and recommendations should be rigorously evaluated experimentally under the local weather conditions.

## Acknowledgments

We wish to thank the extension officers Reuven Birger, Mugira Yunis and Nizar Abed-Al-Ahadi who contributed to this research, Moran Siti, Avida Alon and Dor Rachmani from Luxembourg Industries Ltd. who sprayed the experiments and the growers who allowed us to assess disease development in their olive groves and Bar Ezra Gafniel, who programmed the PedMan system. This research was supported in part by the Israeli Council of Fruit Trees, and the Chief Scientist of Israel Ministry of Agriculture grant #20-02-0039.

## Notes

### Competing Interest Statement

The authors have declared no competing interest.

### Summary of Updates

This updated manuscript incorporates constructive comments from peer reviewers. Revisions include clarifications to the methodology and decision-support workflow, updated figure labeling, and an expanded discussion to better contextualize the findings.

